# Quantitative Restoration of Immune Defense in Old Animals Determined by Naïve Antigen-Specific CD8 T cell Numbers

**DOI:** 10.1101/2021.08.09.453882

**Authors:** Jennifer L. Uhrlaub, Mladen Jergović, Christine M. Bradshaw, Sandip Sonar, Christopher P. Coplen, Jarrod Dudakov, Kristy O. Murray, Marion C. Lanteri, Michael P. Busch, Marcel R. M. van den Brink, Janko Nikolich-Žugich

**Affiliations:** Department of Immunobiology, University of Arizona College of Medicine, Tucson, AZ, USA; University of Arizona Center on Aging, University of Arizona, College of Medicine, Tucson, Tucson, AZ, USA; Program in Immunology, Clinical Research Division, and Immunotherapy Integrated Research Center, Fred Hutchinson Cancer Research Center, Seattle, WA 98109, USA; Department of Immunology, University of Washington, Seattle, WA 98109, USA; Department of Pediatrics, Section of Pediatric Tropical Medicine and National School of Tropical Medicine, Baylor College of Medicine, Houston, TX 77030, USA; William T. Shearer Center for Human Immunobiology, Texas Children’s Hospital, Houston, TX 77030, USA; Blood Systems Research Institute, Vitalant Research Institute, San Francisco, CA, 94118, USA; Department of Medicine and Immunology Program, Memorial Sloan Kettering Cancer Center, New York, NY 10065, USA; Immunology Program, Memorial Sloan Kettering Cancer Center, New York, NY 10065, USA

**Keywords:** Immune aging, immune rejuvenation, IL-7, CD8 T cells

## Abstract

Older humans and animals often exhibit reduced immune responses to infection and vaccination, and this often directly correlates to the numbers and frequency of naïve T (Tn) cells. We found such a correlation between reduced numbers of blood CD8+ Tn cells and severe clinical outcomes of West Nile virus (WNV) in both humans naturally exposed to, and mice experimentally infected with, WNV.

To examine possible causality, we sought to increase the number of CD8 Tn cells by treating C57BL/6 mice with IL-7 complexes (IL-7C, anti-IL-7 mAb bound to IL-7), shown previously to efficiently increase peripheral T cell numbers by homeostatic proliferation. T cells underwent robust expansion following IL-7C administration to old mice increasing the number of total T cells (>four-fold) and NS4b:H-2D^b^-restricted antigen-specific CD8 T cells (two-fold). This improved the numbers of NS4b-specific CD8 T cells detected at the peak of the response against WNV, but not survival of WNV challenge. IL-7C treated old animals also showed no improvement in WNV-specific effector immunity (neutralizing antibody and *in vivo* T cell cytotoxicity). To test quantitative limits to which CD8 Tn cell restoration could improve protective immunity, we transferred graded doses of Ag-specific precursors into old mice and showed that injection of 5,400 (but not of 1,800 or 600) adult naïve WNV-specific CD8 T cells significantly increased survival after WNV. These results set quantitative limits to the level of Tn reconstitution necessary to improve immune defense in older organisms and are discussed in light of targets of immune reconstitution.

## Introduction

Prolonged T cell suppression after myeloablative conditioning is a major clinical problem with considerable clinical implications for therapies such as hematopoietic cell transplantation. Consequently, much work has gone into developing therapies that can boost T cell reconstitution, with varying degrees of success, most of which focus on stimulating thymic function (reviewed in (Granadier, Iovino, Kinsella, & Dudakov, 2021).

Aging is also accompanied by a variable degree of lymphopenia (Groarke & Young, 2019) due to combined effects of early-life thymic involution and late-life defects in lymphocyte maintenance in secondary lymphoid organs (SLO) (rev. in (Nikolich-Žugich, 2018). This lymphopenia in the last third of life correlates to decreased ability of the immune system to respond to infection and vaccination (rev, in (Nikolich-Žugich, 2018). Age-related lymphopenia affects all lymphocytes over time, but is drastically evident for naïve T (Tn) cells that reside in SLO such as the spleen and lymph nodes (LN) (Thompson, Smithey, Surh, & Nikolich-Žugich, 2017). Tn cells are the most diverse T cell subset, in charge of defending the organism against new infection, which transforms Tn cells (CD62LhiCD44lo in mice) specific for the invading infection into effector (Te, CD62LloCD44hi) and subsequently into memory (Tem – effector memory, also CD62LloCD44hi; and Tcm-central memory T cells, CD62LloCD44hiCD49hi). With aging, disturbances in SLO maintenance of Tn cells leads to the accumulation of antigen-independent memory cells (virtual memory; Tvm, CD62LloCD44hiCD49dlo) impacting their phenotype and function and diminishing their ability to protect the organism against infection (Rev. in (Nikolich-Žugich, 2018). It is well established that both CD4 and CD8 Tn cells diminish in numbers by the last third of life in mice (Thompson et al., 2017), non-human primates (Cicin-Sain et al., 2007; Cicin-Sain et al., 2010), and humans ((Groarke & Young, 2019; Wertheimer et al., 2014), albeit with slightly different kinetics depending on the species and compartment sampled. Associations between Tn lymphopenia and decreased immunity in old age were reported. Studies have shown that the CD8 response to live single-cycle poxvirus vaccination directly correlates to the CD8 Tn numbers in non-human primates (Cicin-Sain et al., 2010). Correlations were also found in a study of long-term repeated apheresis subjects, where 15-20% of PBMC were retained by the apheresis machine, and there was a correlation between the number of donations, lymphopenia, and reactivation of persistent herpesviruses (e.g. Varicella-Zoster virus) (Zhao et al., 2021). Moreover, severity of COVID-19 in older adults has been shown to correlate to reduced numbers of CD4 and CD8 Tn cells (Rydyznski Moderbacher et al., 2020). A limitation of these studies is that they did not independently control for aging, and, in most cases, baseline samples were not available to answer whether this was a preexisting condition, a virus-induced lymphodepletion, or a combination of both. All of these intriguing correlative studies raise the possibility of causal association between the (age-related) Tn lymphopenia and functional immunity decline with aging.

Reconstitution of T lymphopoiesis and maintenance in older organisms, also known as immune rejuvenation, has been a highly sought goal in regenerative immunotherapy. Success in that regard has been partial, with no clear translation into an accepted medical protocol. Several treatments in humans and mice transiently (usually up to 45-60 days in mice) reawaken thymic production of Tn cells, including surgical (T. S. P. Heng et al., 2005) and various forms of chemical (Sutherland et al., 2005) sex steroid ablation (SSA), and treatment with various growth factors, including the human growth hormone (Taub, Murphy, & Longo, 2010) and the keratinocyte growth factor (KGF) (Min et al., 2007). Functional evidence that immunity was improved following such treatments in *bona fide* old mice was less abundant (T. S. Heng et al., 2012; Min et al., 2007) and did not involve testing of immune protection using lethal infectious challenge. We have recently discovered that KGF- and pharmacological SSA-induced thymic rebound readily led to restoration, followed by a decline over time, of both thymic cellularity and of the influx of recent thymic emigrants (RTE) into the blood (Thompson et al., 2019) in old mice. Such RTE, however, sparsely populated old spleens and lymph nodes (LN), and treated mice showed no improvement in survival following challenge with West Nile virus (Thompson et al., 2019). That led to investigations of LN involution with age, and its role in immune senescence and reduced immune defense with aging.

As part of these studies, we sought to conclusively address the role of age-linked lymphopenia in reduced immune defense with aging. Here, we report using IL-7 complexes (IL-7C, IL-7 bound to an anti-IL-7 Ab, (Boyman, Ramsey, Kim, Sprent, & Surh, 2008) to increase lymph node cellularity by means of homeostatic proliferation and found a doubling of CD8 Tn precursors specific for the immunodominant NS4b epitope of WNV. However, most of these cells were no longer true naïve CD8 cells, but rather antigen-independent Tvm. Upon challenge with WNV, we found no improvement of immune protection, and no increase in either CD8 effector T cells or their cytotoxic function, consistent with our prior findings that Tvm cells do not proliferate well upon stimulation. These mice also did not produce robust anti-WNV antibodies, suggesting that CD4 and/or B cell defects were not remedied either. To evaluate whether an adequate reconstitution of true naïve cells could protect old mice against lethal challenge, we transferred graded amounts of naïve CD8+ NS4b/Db-specific TCR transgenic cells. We found that numbers of Ag-specific Tn cells needed to confer protection to old mice were slightly above the numbers of Ag-specific Tn precursors present in young adult mice, suggesting that both the numbers and the quality of precursors are important in reconstitution. These results establish the numerical limits of CD8 Tn reconstitution and are discussed in light of strategies for immune reconstitution with aging.

## RESULTS

### Older age and reduced naïve T cell numbers are associated with WNV disease severity

Previously unencountered and emerging viral infections often cause increased disease severity and mortality in older populations. Indeed, patients who have experienced the most severe symptoms caused by WNV were significantly older than those who had milder disease ((Campbell, Marfin, Lanciotti, & Gubler, 2002) and Figure 1A), and similar findings have been made in other emerging infectious diseases including SARS-CoV, Chikungunya virus, and SARS-CoV-2. Consistent with our prior data in general geriatric populations (Wertheimer et al., 2014) our cohort of WNV-exposed participants exhibited significantly lower levels of circulating CD8, but not CD4, Tn cells when compared individuals who experienced asymptomatic infection (Figure 1B, C). We determined T cell phenotype frequencies using a combination of CCR7, CD45RA, CD28 and CD95, to define naïve T cells as CCR7hi,CD45RAhi,CD28int/hi and CD95low. Participants who experienced the most severe symptoms, meningitis and encephalitis, exhibited a measurable loss of both Tn CD4 and CD8 T cells as compared to at least one other group. In a retrospective study, where a limited number of patient samples were collected long after asymptomatic infection or disease recovery, it is impossible to determine whether this was a cause, an epiphenomenon, or an effect, of their severe infection. To examine the nature of associations between age, Tn cell deficit, and severe WNV disease, we examined this issue in an established mouse model of age-related susceptibility to WNV (Brien, Uhrlaub, Hirsch, Wiley, & Nikolich-Zugich, 2009).

**Figure 1.**
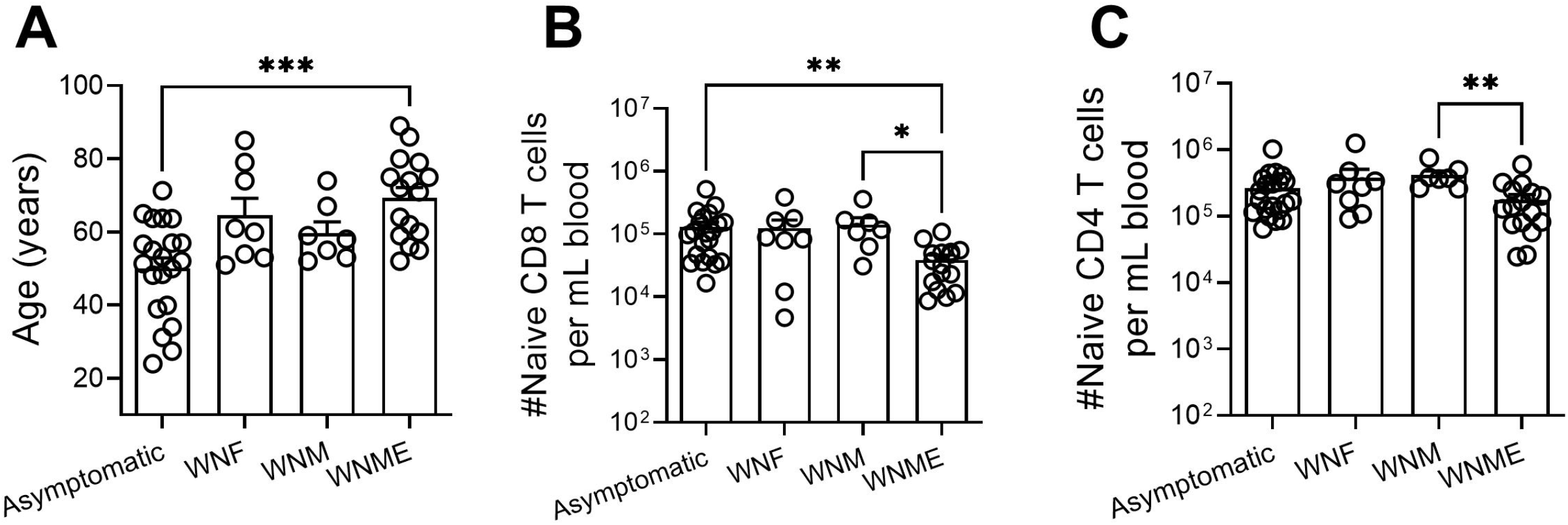
Older adults experience more severe WNV disease and have a dearth of naïve CD8 T cells. A cohort of 52 patients with WNV disease severity clinically classified as Asymptomatic, WN Fever (WNF), WN Meningitis (WNM), and WN meningitis and encephalitis (WNME) were retrospectively recruited to participate in our study. The age of the patients experiencing the most severe disease is significantly older than asymptomatic (A). Naïve (CD95^low^ and ^CD28int/high^, also confirmed additionally to be CCR7hi and CD45RA+) CD4 and CD8 T cells were phenotyped by FCM and counts were extrapolated from clinical cell blood count (CBC) data collected on the fresh blood sample. Statistical significance evaluated by Kruskal-Wallis with Dunn’s multiple comparisons test between all groups, +SEM. \**P* ≤ 0.05; \*\**P* ≤ 0.01; \*\*\**P* ≤ 0.001.

### The murine model recapitulates the descriptive findings from human subjects

It is well established that the naïve T cell compartment contracts with age and is proportionally less well represented than the T cells carrying memory phenotypes (Zhang, Weyand, & Goronzy, 2021). A direct correlation with disease severity, however, has not been conclusively established. To determine if our WNV mouse model is valid to interrogate this association, we measured the number of circulating CD8 Tn cells in WNV-infected adult (3-5 month old) and old (19-22 month old) mice on day 7 p.i.; a time point prior to onset of WNV disease (Figure 2A). Mice that will eventually succumb to WNV exhibited lower numbers of circulating naïve CD8 and CD4 T cells at this early time point when compared to those that will survive (Figure 2B, C). This disparity held regardless of age, despite the fact that old mice exhibit reduced numbers of Tn cells overall (Figure 2B, C). These data support the possibility that a numeric advantage in naïve T cells is directly linked to WNV resistance regardless of age. These results are also consistent with the human data from Figure 1 with the exception that there is also a robust loss of naïve CD4 T cells in susceptible mice. Of note, there was overlap in CD8 Tn numbers between humans with severe and asymptomatic WNV disease, as well as between mice that succumbed and survived WNV infection. This suggests that the numbers of CD8 Tn cells are not the only determinant of increased WNV resistance, consistent with the extensive literature on the multipronged nature of immune defense against WNV (Brien et al., 2009; Purtha et al., 2007; Richner et al., 2015). The overlap in the human and mouse data provides confidence that this mouse model can be used to examine whether, and to what extent, reversing age-induced lymphopenia and, specifically, a deficit in Tn cells, may afford improved immune protection against WNV.

**Figure 2.**
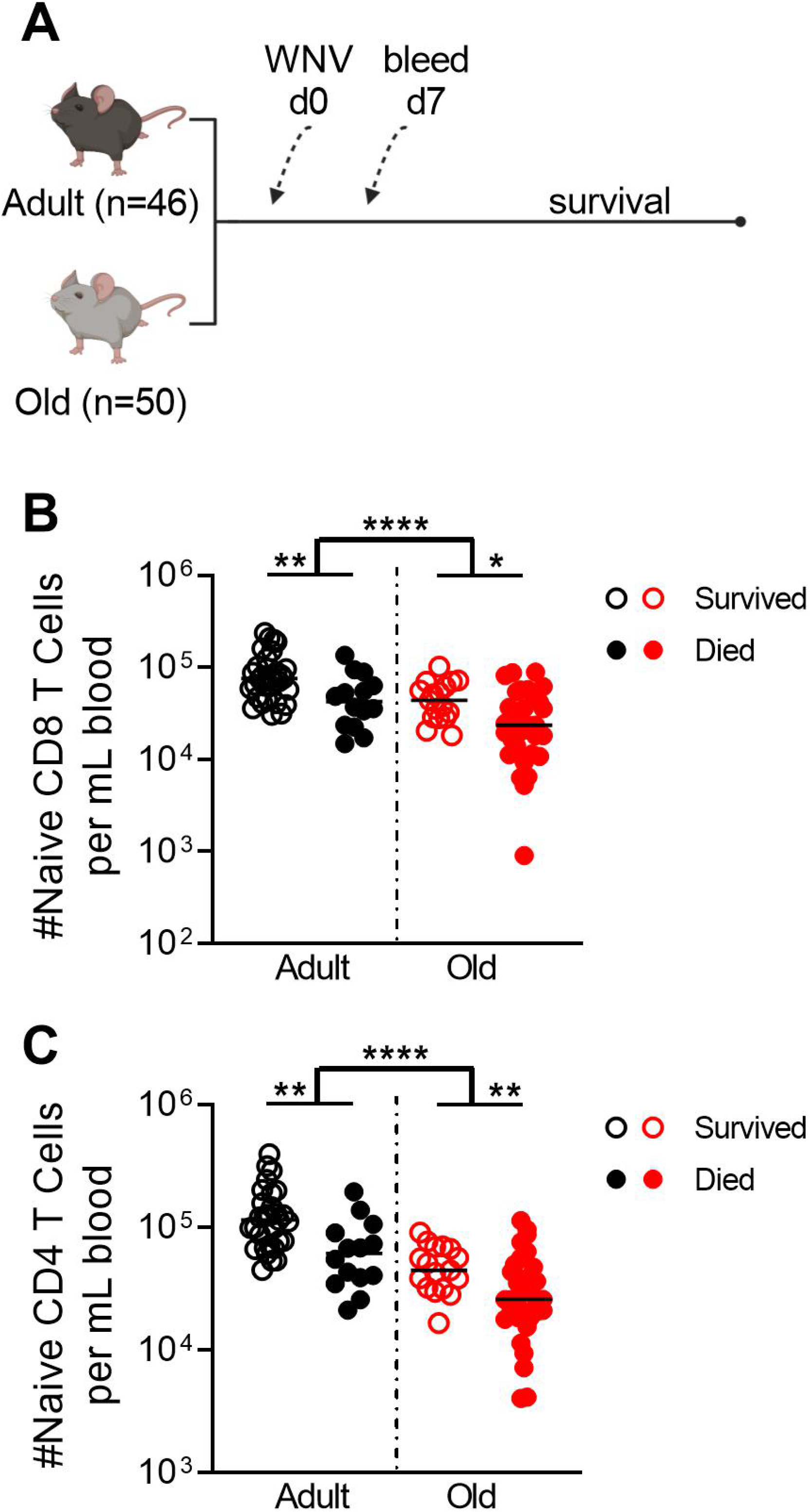
Decreased numbers of naïve CD8 T cells in both young and old mice correlate with increased susceptibility to WNV. Adult (3-5 months) and old (19-22 months) C57BL/6 mice were infected with WNV, 1000pfu, f.p.; bled on day 7 for FCM; and followed for susceptibility (A, schema). Absolute numbers of naïve CD8 or CD4 T cells were enumerated per mL of blood on day 7 post-infection by FCM (B, C). Data compiled from two independent experiments. Statistical significance evaluated by t-test, median. \**P* ≤ 0.05; \*\**P* ≤ 0.01; \*\*\*\**P* ≤ 0.0001.

### IL-7 complex treatment increases the number of circulating Tn cells

IL-7 is a pleiotropic cytokine that provides an indispensable homeostatic survival signal to Tn cells, and that drives Tn and other T cell subsets into homeostatic proliferation under the conditions of lymphopenia. Delivered *in vivo* as a cytokine-antibody complex, IL-7C has been shown to drive naïve T cell proliferation and increase thymopoeisis in unmanipulated as well as lymphopenic mice (Boyman, Kovar, Rubinstein, Surh, & Sprent, 2006). Despite its therapeutic impact in adult mice, this model has not been rigorously interrogated for reversal of age-related lymphopenia and reduced susceptibility to infection. We treated old mice with IL-7C (3 i.p. injections, every other day), and sacrificed them on days 7, 10, and 14 post-first IL-7C treatment to assess total spleen and LN content (schema, Figure 3A). Total cellularity of secondary lymphoid tissues (spleen and LNs) remained elevated through day 14 post-first treatment (Figure 3B), an expansion mirrored by the numbers of CD4 and CD8 T cells (Figure 3C, D). Additionally, numbers of CD4 and CD8 Tn cells in treated old mice were no longer statistically significantly reduced relative to those in adult mice (Figure 3E, F). The numerical increase in CD4 T cells was driven by split increases in Tcm and Tvm phenotype cells, neither of which were independently statistically significant (Figure 3G, H). In the CD8 pool, T cells with a CM or Tvm phenotype dominated the increase in numbers (Figure 3I, J). To determine whether the expansion of T cells by IL-7C also included CD8 T cells specific for the immunodominant epitope of WNV, NS4b_2488_, we enumerated the number of these CD8 T cell precursors on day 13 post-treatment in unimmunized mice (Figure 3K). Similar to our prior work with other immunodominant CD8 epitopes (Rudd, Venturi, Davenport, & Nikolich-Zugich, 2011; Smithey, Li, Venturi, Davenport, & Nikolich-Žugich, 2012), we found a marked ~62% loss of NS4b-specific CD8 precursors in old animals compared to adult counterparts (Figure 3F). We found that IL-7C treatment led to a numerical increase in CD8 precursors specific for the WNV NS4b immunodominant epitope that, on average, no longer statistically differed from that in adult animals (Figure 3K). The response between individual animals, however, was heterogenous, with many animals exhibiting no increase, and others exhibiting up to a two-fold increase in precursor numbers. Further, and as would be expected, NS4b-specific precursors were of a similar phenotype as compared to the rest of the CD8 T cell pool (Table 1).

**Figure 3.**
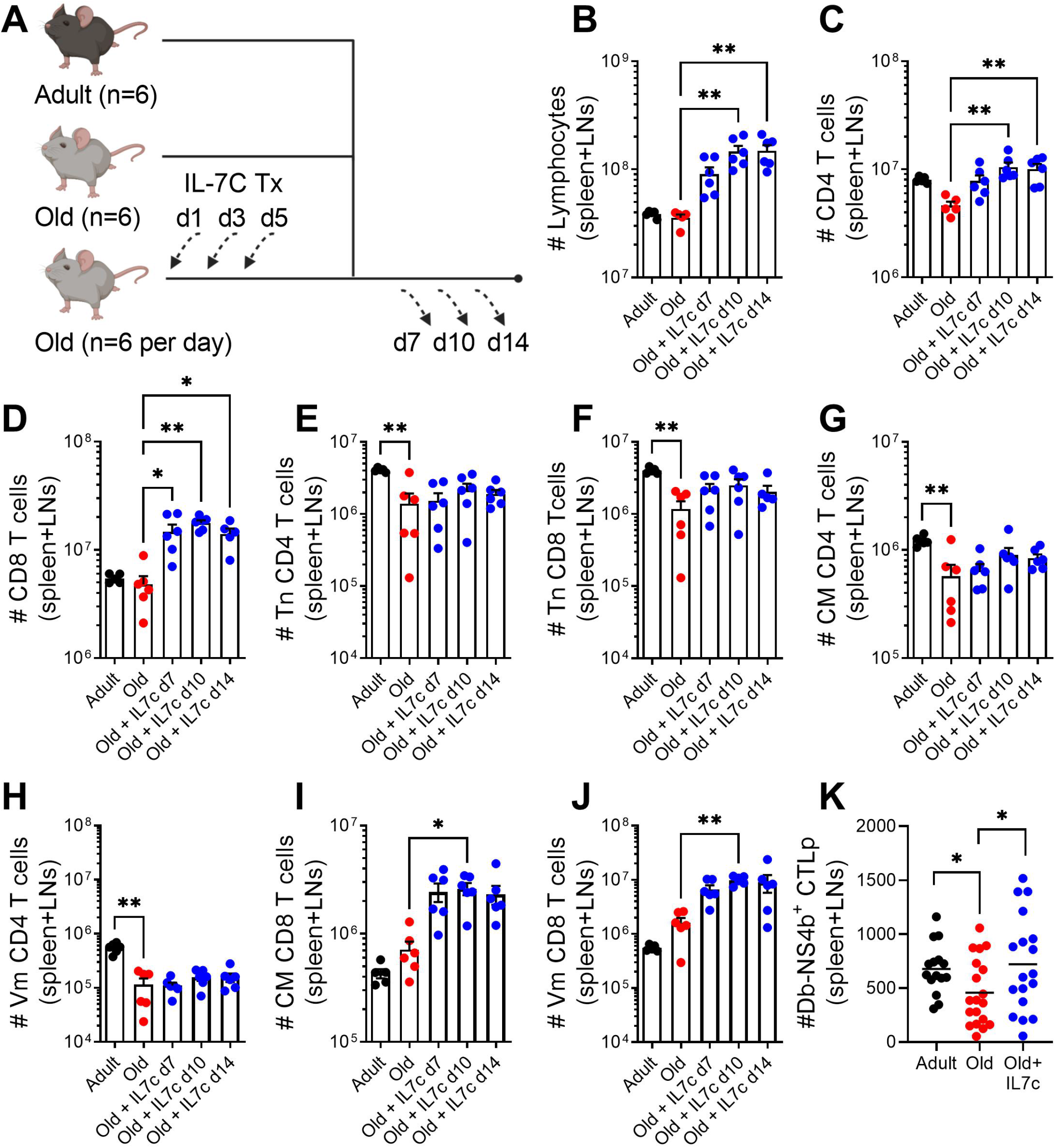
IL-7C treatment expands the total number of T cells available in secondary lymphoid tissues. Old (19-22 months) C57BL/6 mice were treated with IL-7C on days 1, 3, and 5 of the experiment and sacrificed on days 7, 10, and 14 post-first treatment alongside untreated adult (3-5 months) and old (19-22 months) (schema, A). The total number of lymphocytes (B), CD4 T cells (C), and CD8 T cells (D) are increased at indicated day(s) post-first treatment. The total number of naïve, CD44 low and CD62L high, CD4 and CD8 T cells are not significantly increased at any day post-treatment (E, F). The expansion of CD4 T cells is not dominated by either CM (CD44 high, CD62L high, and CD49d high) or virtual memory (CD44 high, CD62L high, and CD49d low) phenotyped cells (G,H). Both CM and VM phenotype CD8 T cells are expanded at day 10 post-first treatment (I,J). WNV-specific CD8 CTLp (Db-NS4b_2488_) are not reliably increased by IL-7C treatment (K). (B-J) Data shown is representative of two experiments with similar results and statistical significance evaluated by one-way Anova with Dunn’s multiple comparisons test. All comparisons are to untreated old mice. \**P* ≤ 0.05; \*\**P* ≤ 0.01. (K) IL-7C treated group is day 13 post-first treatment. Data compiled from three independent experiments and statistical significance evaluated by t-test comparison to O, median. \**P* ≤ 0.05.

**Table 1.**

| Table 1 |  | Adult | Old | Old+IL7-C |
| --- | --- | --- | --- | --- |
| Tn | %CD44<sup>low</sup> CD62L<sup>high</sup> of Db-NS4b CTLp (+/-SEM) | 65.1<br>(2.6) | 56.2<br>(4.2) | 47.6<br>(7.1) |
| Tvm | %CD49d<sup>low</sup> of CM Db-NS4b+ CTLp (+/-SEM) | 93.7<br>(2.0) | 75.7<br>(6.8) | 83.8<br>(4.5) |

The relative improvement in total T cell numbers, from days 10 to 14, including an overall increase in WNV-specific CD8 precursors on day 13 following IL-7C treatment suggested that functional immunity could be improved within this temporal window, because lymphopenia in old mice was measurably reduced and residual IL-7C (if any) did not cause further T cell proliferation in secondary lymphoid tissues.

### IL-7c treated old mice gain numbers, but not function, of WNV-specific CD8 T cell responses

To conclusively test whether this increase in T cell numbers is sufficient to overcome infectious disease susceptibility, we challenged IL-7C treated mice with WNV. Based on the above kinetics of IL-7C action, we chose day 13 as an optimal time point for challenge, reasoning that the mice would benefit from an expanded T cell pool at the time when IL-7C is no longer driving further proliferation of T cells, and should not influence the development of a functional immune response to WNV. Adult (3-4 months); old (19-22 months); and IL-7C treated old (19-22 months) mice were challenged with 1000pfu of WNV and followed for WNV-specific CD8 T cell responses and survival (Figure 4A). On day 7 post-infection, old mice treated with IL-7C showed a 3-fold improvement in the number of WNV NS4b-specific CD8 T cells measured in the blood (Figure 4B). However, this improvement failed to translate to better survival (Figure 4C), despite the fact that we observed roughly equivalent numbers of WNV-specific CD8 T cells in old IL-7C treated mice compared to adult unmanipulated mice. When we segregated each experimental group into animals that died or survived WNV challenge, we found that those that survived again exhibited a numeric advantage in WNV-specific cells over those that succumbed (Figure 4D). The effect, however, was not robust enough to ensure that the group, as a whole, survives better than their untreated counterparts.

**Figure 4.**
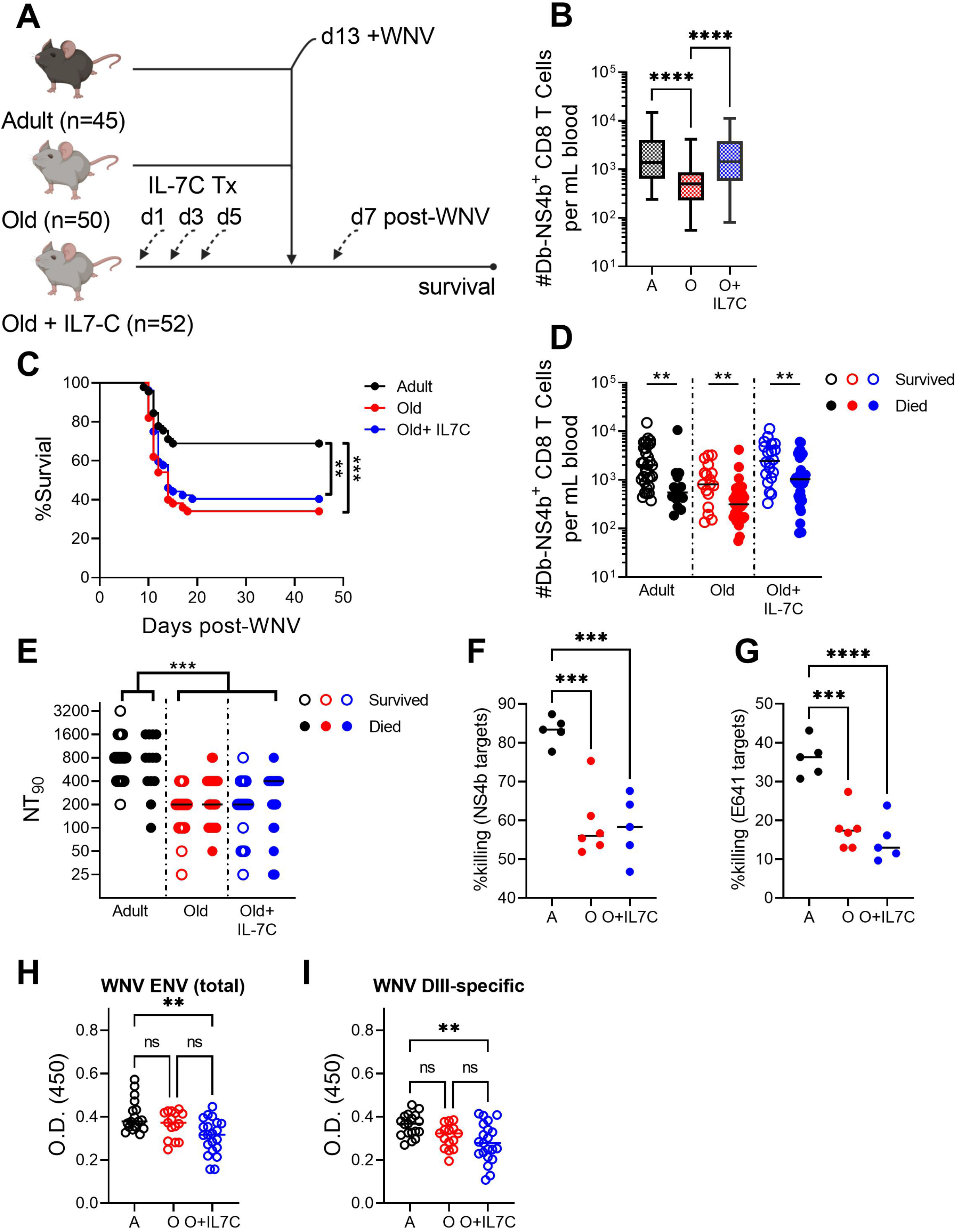
IL-7C treatment of old mice improves WNV-specific CD8 T cell responses without impacting survival. Adult (3-5 months), Old (19-22 months), and Old (19-22 months) mice treated with IL-7C 13 days before challenge are infected with WNV (1000pfu, f.p.; n=45-52 per group). On day 7 post-WNV infection mice are bled and monitored for survival (schema, A). Number of WNV-specific CD8 T cells per mL of blood on day 7 post-WNV (B). WNV survival at day 45 post-infection (C). Number of WNV-specific CD8 T cells from (B) broken by those that survived and those that died from WNV in each group (C). Neutralizing antibody titers on day 7 post-WNV broken by those that survived and those that from died from WNV in each group (E). *In vivo* killing assay on day 7 post-WNV for NS4b_2488_ (F) and ENV_641_ (G). Total WNV-ENV and DIII-specific memory antibody levels >45 days post-infection (Figure H, I). Data compiled from two experiments. Statistical significance evaluated by anova in (B, F, G) and t-test in (D) and (E) as represented on graphs. Significance of survival evaluated by Log-rank (Mantel-Cox) test. \**P* ≤ 0.05; \*\**P* ≤ 0.01; \*\*\**P* ≤ 0.001; \*\*\*\**P* ≤ 0.0001.

Multiple effector mechanisms of adaptive immunity must synergize to clear WNV infection. These include potent neutralizing antibodies as well as CD4 and CD8 cytotoxic T cell effector function. To mechanistically determine whether any functional improvement in immunity was restored by IL-7C treatment we interrogated the immune response in IL7-C treated old mice in a step-wise manner. IL-7C have the potential to enhance B cell lymphopoiesis, and to thereby increase B cell numbers, although they are not expected to directly affect the mature B cell compartment (Boyman et al., 2008). Previous studies have reported that IL-7C co-delivered with immunization can boost humoral immune responses through enhanced generation of germinal center B cells and Tfh cells (Seo et al., 2014). Therefore, we first analyzed neutralizing antibody titers at day 7 post-WNV infection as a proxy for generalized improvement in B cell responses. We detected no improvement in WNV neutralization by sera of old mice treated with IL7-C relative to untreated controls (Figure 4E). We next examined whether WNV-specific CD8 or CD4 T cells exhibited improved killing function following IL-7C treatment by measuring cytotoxicity in vivo (Figure 4F, G). We included targets specific for one WNV-specific CD4 immunodominant epitope, ENV_641_, in this assay since CD4 T cells are known to mount cytotoxic responses against WNV in adult mice and data about their specific contribution to viral clearance with age is limited (Brien, Uhrlaub, & Nikolich-Zugich, 2008). We found that, despite numeric increases in circulating NS4b-specific CD8 T cells, IL-7C treated old mice did not have improved cytotoxic capacity to clear NS4b_2488_ peptide-loaded targets as compared to untreated old mice (Figure 4F). There was also no improvement in killing of the ENV_641_-loaded targets (Figure 4G), consistent with the fact that we saw no increase in CD4 Tn cell numbers (Fig. 3E). These data indicate an overall lack of improvement in cytotoxic function in old mice treated with IL-7C. As these assessments were done prior to the onset of disease, we measured memory antibody responses in WNV survivors as an indication of whether IL7-C treatment impacted overall humoral immunity. We determined that neither total envelope nor DIII lateral ridge specific antibodies were improved by IL7-C treatment prior to WNV infection (Figure H, I). In fact, when comparing the survivors of WNV infection, it appears that old mice treated with IL-7C established an overall lower level of circulating WNV-specific antibodies at this point in memory (Figure H, I). Our interpretation of this data is that the expansion of CD4 T cells by IL7-C treatment does not afford a short- or lasting-impact on antibody responses to WNV infection. At face value, these results suggest that numerical improvement in CD8 and CD4 T cells afforded by IL-7C treatment is not sufficient to improve functional immunity.

To investigate why a numeric increase in antigen-specific precursors was not able to restore T cell responses we considered the possibility that the expansion induced via IL7-C treatment may push the cells into an exhausted state, impairing their ability to participate in the WNV response. T cells with an ‘exhausted’ phenotype are typically seen in the course of chronic infections and cancer (Collier et al., 2021; Reiner, 2021). The number of divisions and other factors required to induce this state are not fully understood. To investigate whether homeostatic proliferation driven by IL-7C treatment increases the proportion of responding WNV-specific T cells with an exhausted phenotype we measured exhaustion markers in both adult and old mice treated with IL7-C on day 7 of WNV. We included an adult cohort in these experiments to dissociate aging from treatment-associated phenotypes. Our data confirm that a higher percentage of WNV-specific CD8 T cells in aged mice express PD-1, LAG-3, or 2B4 regardless of whether they have been treated with IL7-C (Figure 5A, B, C). The expression of these molecules could contribute to their poor cytotoxic function, and overall lack of improvement in survival, experienced by the old animals. These data suggest that the homeostatic proliferation induced by IL7-C treatment is not *per se* pushing T cells into an exhausted state. Rather, consistent with some reports (Decman et al., 2012) we hypothesize that the age of the animal may have already determined the phenotype of the available T cell pool and this state is maintained through the expansion initiated by IL7-C treatment to deleterious effect.

**Figure 5.**
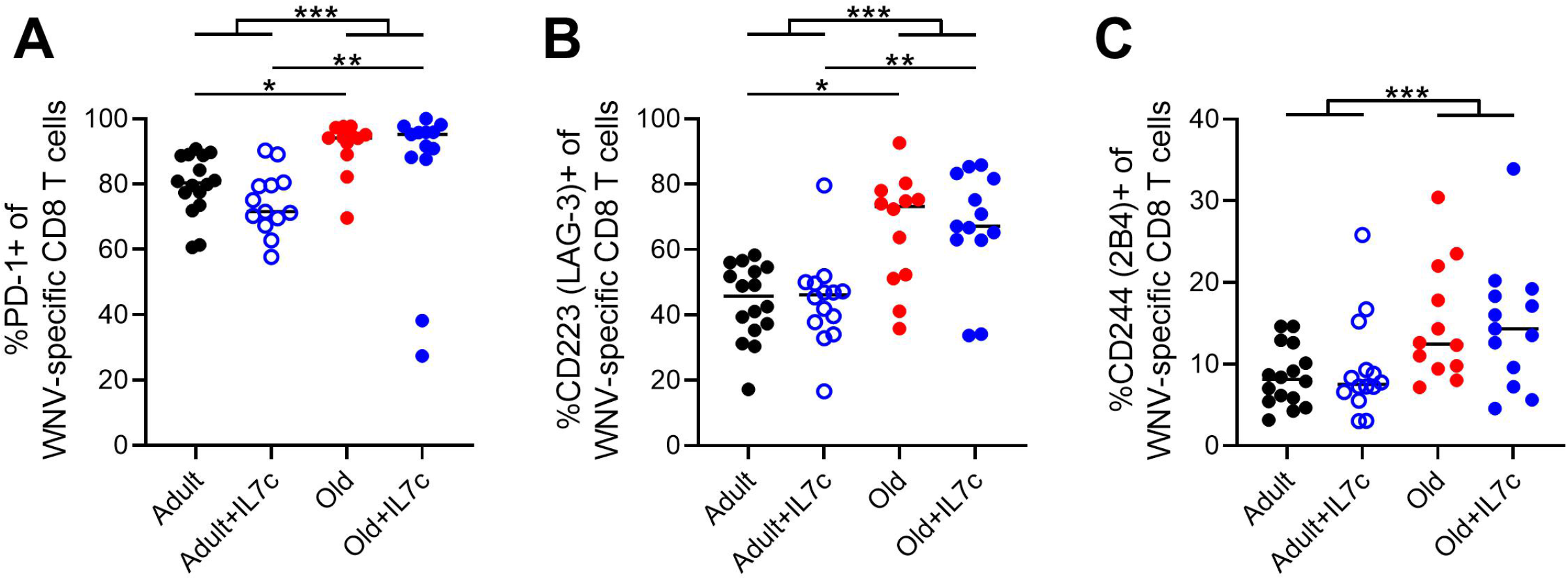
IL7-C treatment does not induce T cell exhaustion. Adult (3-5 months) and Old (19-22 months) mice were treated with IL-7C 13 days before infection with WNV (1000pfu, f.p.). On day 7 post-WNV infection mice were bled and circulating PBMC’s were analyzed by FCM for expression of PD1, CD244 (2B4), and CD223 (LAG-3) on NS4b_2488_+ CD8 T cells (A-C). Data compiled from two experiments. Statistical significance evaluated by anova in (all groups) and t-test in (by age) as represented on graphs. \**P* ≤ 0.05; \*\**P* ≤ 0.01; \*\*\**P* ≤ 0.001.

However, the fact that a similar increase in precursor numbers induced by IL-7C treatment did not lead to either improved immunity or improved cytotoxicity, suggested that some other property of reconstituted cells may be of critical importance. To directly address whether an increase in CD8 Tn precursors specific for an invading pathogen can overcome the age-related defects in immune defense and afford improved survival to old mice, we performed adoptive transfer experiments. We transferred graded levels of WNV-specific CD8 T cells (NS4b_2488_ specific, referred to as WNV-I (Aguilar-Valenzuela et al., 2018)) the day before challenge into old mice to boost the number of precursors available to respond. These transgenic T cells have a naïve phenotype at time of transfer and are not yet primed or cytotoxic. After challenge, the transferred cells quantitatively restored the WNV-specific CD8 T cell numbers measured in the blood on day 7 post-infection (Figure 6A). The differences between old mice treated with IL-7C and those receiving adoptively transferred cells was that the former mounted a response from endogenous, old precursors, and these precursors were driven into homeostatic proliferation by IL-7C; the latter received naïve, unperturbed NS4b-specific precursors from adult donors. Both the age and the immediate prior proliferation induced by IL-7C could have impacted their response. More importantly, in the transfer experiments, we were able to significantly improve survival in old mice following WNV challenge with 5,400 transferred WNV-specific CD8+ Tn cells; but not with the two lower transfer boluses of 1,800 and 600 cells (Figure 6B).

**Figure 6.**
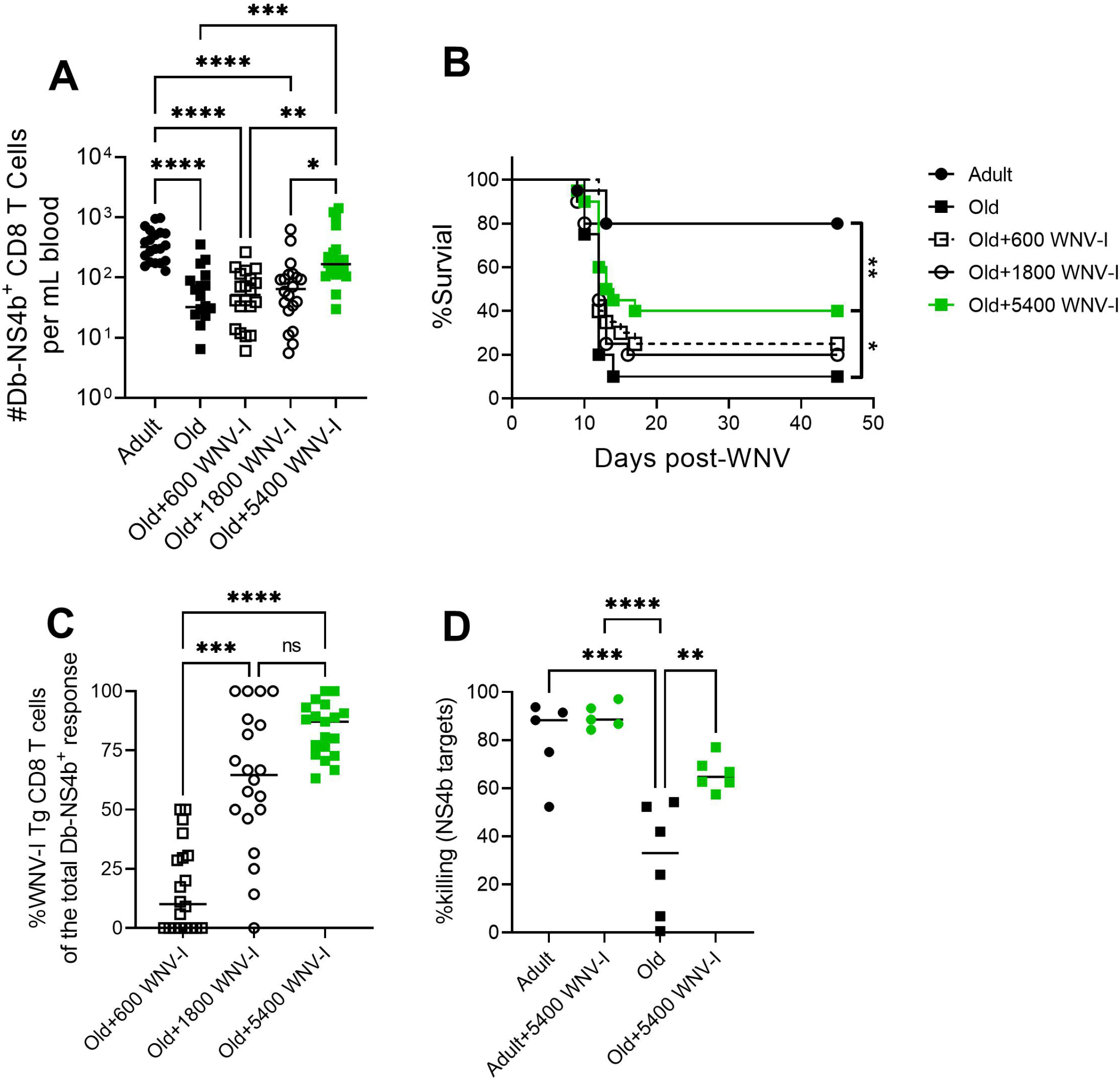
Adoptive transfer of WNV-I Tg CD8 T cells rescues old mice from WNV susceptibility. WNV-I CD8 T cells (5400, 1600, or 600) were transferred to groups of old mice (19-22 months) alongside adult (3-5 months) and Old (19-22months) mice the day before WNV challenge (1000pfu, f.p.). On day 7 post-infection mice were bled and total numbers of circulating WNV-specific CD8 T cells were enumerated per mL of blood (A). Mice were monitored for survival to day 45 post-infection (B). The frequency of donor WNV-I CD8 T cells (CD45.1) contributing to the response in (A) is shown in (C). *In vivo* killing assay on day 7 post-WNV for NS4b_2488_ (D). Data compiled (A-C) or representative (D) of two experiments. Statistical significance evaluated by anova (A, C, D) with multiple comparisons test. Significance of survival evaluated by Log-rank (Mantel-Cox) test. \**P* ≤ 0.05; \*\**P* ≤ 0.01; \*\*\**P* ≤ 0.001; \*\*\*\**P* ≤ 0.0001.

To put these numbers in the context of physiological precursor numbers available to old and adult animals, we determined the efficacy of engraftment of transferred cells. Some estimates of transfer efficiency posit that only about 10% of the transferred T cells engraft in i.v. injected mice (Blattman et al., 2002; Moon et al., 2009). We enumerated the total number of NS4b-specific CD8 T cells that were either endogenous (CD45.2) or transgenic (CD45.1). Indeed, our experiments confirmed that 24h after transfer, we could recover between 12-15% of transgenic cells (Supp. Fig 1A). Note here that the numbers of engrafted cells were quantified in direct comparison to the number of endogenous naïve precursors recovered from adult or old mice as they were enumerated alongside each other (Supp. Fig 2). This means that a dose of 5,400 transferred cells reconstituted about 650-810 NS4b precursors, that were effectively capable to significantly protect the old animal against WNV with survival not significantly different from untreated adult mice. Of interest, an adult mouse contains 500-700 NS4b-specific CD8+ precursors. Importantly, old mice that received 5400 WNV-I Tg CD8 T cells efficiently recruited them to the WNV response (Figure 6C) and exhibited improved cytotoxic function measured by increased killing (Figure 6D). This offers a plausible mechanism for improved survival of this group (Figure 6B). We conclude that restoring the precursor numbers of an immunodominant CD8 Tn specificity to adult levels, and with cells that have a Tn phenotype, may be enough to confer a significant degree of protection to an old mouse against WNV, despite concurrent defects in many other arms of the immune response.

## DISCUSSION

Immune therapies that seek to overcome age-induced lymphopenia are an attractive strategy to reduce susceptibility to infectious diseases in older adults. Conceptually, increasing the total cellular resources of the immune system, regardless of antigen specificity, would provide the best solution for broad-based protection against diverse challenges. IL-7C treatment has been shown to increase peripheral T cell numbers even in the absence of thymic output (Boyman et al., 2008). In this work, we formally demonstrated that IL-7C treatment can transiently increase T cell numbers in old mice that have suffered substantial thymic involution, predominantly due to an expansion of virtual central memory Tvm phenotype cells. We found significant, but variable, expansion of the numbers of CD8 precursors specific for WNV, and an overall increase in both CD8 and CD4 T cell numbers. However, these expanded cells did not protect mice from WNV infection, and we could find no evidence that the cytotoxic function of either CD8 or CD4 T cells was improved in IL-7C treated old mice. While we did not enumerate B cells, their protective responses were similarly flat, with no improvement in neutralization capacity, despite an increase in total numbers of CD4 T cells. Therefore, under our experimental conditions, IL-7C treatment showed no appreciable immune protection benefit to the old mice, similar to our prior studies with unconjugated IL-7 in non-human primates (Okoye et al., 2015).

Given that protection against WNV normally requires a coordinated and multi-faceted response, it may not be surprising that the increase in T cells alone cannot augment protection. It was more puzzling, however, that we could find no evidence of improved responses from the numerically increased T cell pool. To further explore this question and test whether we could improve protection against WNV by boosting a single arm of protective immunity, we performed adoptive transfer of graded numbers of transgenic CD8 T cells. We demonstrated that sufficient numbers of transferred cells (approximately equivalent to the precursor content of a single adult mouse) were able to significantly improve survival, albeit not to restore it to the levels of adult mice. These results begin to provide quantitative targets of CD8 T cell reconstitution, and also provide a proof-of-principle that the aged immune system can coordinate an effective immune response if the proper T cell resources are provided.

One key unanswered question remained regarding the relationship between precursor T cell numbers and their quality. Specifically, adoptively transferred old mice contained similar numbers of overall precursors relative to the IL-7C treated old mice, suggesting that somehow the quality of the two precursor populations was different. To address the potential differences, we considered prior work from ours and other groups on age-related changes in Tn cell maintenance. Specifically, with aging, an increasing number of Tn cells change their maintenance pattern and become Tvm (Chiu, Martin, Stolberg, & Chensue, 2013; Decman et al., 2012; Rudd et al., 2011) to the point that most of the central memory phenotype cells in SPF mice are of Tvm origin. While work from other groups (Lee, Hamilton, Akue, Hogquist, & Jameson, 2013) and us (Renkema, Li, Wu, Smithey, & Nikolich-Žugich, 2014) demonstrated that Tvm cells in young adult mice exhibit strong proliferation and enhanced effector function relative to Tn cells, we have also shown that with age Tvm cells exhibit reduced proliferative response to cognate antigen (Renkema et al., 2014). By contrast, the response to homeostatic cytokines remains preserved in Tvm cells. Additional studies will be necessary to directly compare in vivo antiviral function and protective capacity of Tvm and Tn cells of different ages with and without prior expansion by IL-7C and similar treatments. Moreover, as these studies were focused on improving immune protection against acute lethal challenge, it remains to be investigated whether and to what extent the interventions used here, including IL-7C, could potentially improve responses to vaccination, including memory responses. That notwithstanding, the work presented here places quantitative limits on desirable levels of immune reconstitution and opens provocative questions about their quality, that remain to be answered in the future.

## EXPERIMENTAL PROCEDURES

### Study Subjects and Blood Samples

This study was approved by the Institutional Review Boards at the University of Arizona (Tucson, AZ; # 080000673; currently #2102460536), the Oregon Health and Science University (Portland, OR; # IRB00003007); the University of Texas Health Science Center at Houston, TX; and the Blood Systems Research Institute, San Francisco, CA. - presently Vitalant Research Institute. Exclusion criteria included known immunosuppressive pathology, stroke, cancer, or use of steroids within the last 5 years. WNV-infected donors were enrolled by the University of Texas at Houston, 2006–2009 or Blood Systems Research Institute (BSRI), 2009 – 2011. Samples, demographics (Supp. Table I), and symptoms data were collected and analyzed as previously described (Lanteri et al., 2014) after the subjects provided an informed consent approved by the UCSF Committee on Human Research (protocol #10-01255) or the Committee for the Protection of Human Subjects at the University of Texas Health Science Center at Houston (HSC-SPH-03-039), respectively. Blood was drawn into heparinized Vacutainer CPT tubes (BD Bioscience) and processed fresh at respective sites per manufacturer’s recommendations to isolate peripheral blood mononuclear cells (PBMC) and plasma; K2-EDTA tubes were used to determine complete blood counts. PBMCs were frozen in 90% fetal bovine serum (FBS) and 10% DMSO.

### Flow Cytometry – human samples

Frozen PBMCs were thawed in RPMI medium supplemented with 10% FBS, penicillin and streptomycin in the presence of DNAse (Millipore Sigma) then rested overnight in X-Vivo medium (Lonza) supplemented with 5% human male AB serum. 1–3 ×10^6^ PBMC were stained with a saturating concentration of the following antibodies: CD3 AF700 (clone SP34-2, BD Biosciences), CD4 eFluor450 (clone OKT4, eBioscience), CD8β ECD (2ST8.5H7, Beckman Coulter), CD95 APC (clone DX2, BD Biosciences), and CD28 PercpCy5.5 (clone CD28.2, BioLegend) in 1XPBS, 2% FBS and next incubated with LIVE/DEAD® Fixable Dead Cell Stain (ThermoFisher Sci), both for 30 min at 4°C. Flow cytometry acquisition was performed on a BD Bioscience Fortessa and analyzed using FlowJo software (Tree Star).

### Mice

Mouse experiments were carried out in strict accordance with the recommendations in the Guide for the Care and Use of Laboratory Animals of the National Institutes of Health. Protocols were approved by the Institutional Animal Care and Use Committee at the University of Arizona (IACUC 19-580, PHS Assurance No. A-3248-01. Old (19-22 months) and adult (3-5 months) were obtained from the National Institute on Aging Rodent Resource or Jax laboratories. Mice in each experiment were age matched within 4 weeks. WNV-I mice were obtained from Drs Jason Netland and. Michael J. Bevan (Aguilar-Valenzuela et al., 2018). Footpad (f.p.) injections and retro-orbital intravenous cell transfers were performed under isoflurane anesthesia. IL-7 cytokine was obtained from the Biological Resources Branch (BRB), Developmental Therapeutics Program (DTP), Division of Cancer Treatment and Diagnosis (DCTD), National Cancer Institute (NCI). IL-7R antibody (clone M25) was purchased from BioXcell. 0.59ug of IL-7 was complexed with 2.9ug of M25 antibody for 30 min at RT then administered via intraperitoneal injection in 200uL USP saline on days 1, 3, and 5 of the experiment. Euthanasia was performed by isoflurane overdose or cervical dislocation.

### Virus

West Nile virus NY 385–99, isolated from the liver of a snowy owl, and in its second passage, was a kind gift from Dr. Robert Tesh (University of Texas Medical Branch at Galveston, Galveston, TX). Virus was prepared as previously described (Brien et al., 2009). Mice were infected by footpad (f.p.) route with 1,000 pfu WNV/50 μl/mouse and monitored for survival until 45 days p.i.

### Tissue harvest, cell counts, flow cytometry, neutralization titers, antibody ELISA

Blood was collected via retro-orbital route and subjected to hypotonic lysis of red blood cells. Spleen and peripheral lymph nodes (inguinal, brachial, axillary, and cervical) were collected and disassociated over 40uM mesh. Cells were resuspended and stained overnight at 4°C in a saturating concentration of antibodies against CD49d (9C10, MFR4.B), CD44 (IM7), CD62L (MEL-14), CD3 (17A2), CD4 (RM4–5), CD8a (53–6.7), CD45.1 (A20), CD244.2 (m2B4 (B6) 458.1), CD223 (LAG-3, C9B7W), PD-1 (29F.1A12) (eBioscience and BioLegend) and H2-Db/NS4b tetramers (NIH Tetramer Facility, Atlanta, GA) followed by Live/Dead viability dye (ThermoFisher Sci). Cytotoxic precursor enumeration was accomplished as described in (Uhrlaub, Smithey, & Nikolich-Žugich, 2017). NS4b-Tg CD45.1 CD8 T cells were purified with Miltenyi Biotec reagents and columns using negative magnetic selection. Cell counts and purity (>95%) were confirmed by FCM analysis and count beads before adoptive transfer. Optimization procedures for flow cytometry are based on current standards and included full-minus-one (FMO) gating controls. Samples were acquired on a BD LSRFortessa cytometer (BD Biosciences) or Cytek Aurora (Cytek Biosciences) and analyzed with FlowJo software (Tree Star). WNV neutralization titers were performed as described in (Wong et al., 2020) and scored as a 90% reduction in viral inoculum. WNV ELISA’s for both total ENV and DIII were also performed as described in (Wong et al., 2020) and directly reported as O.D.

### In Vivo Killing Assay

Splenocytes from C57BL/6 mice were disassociated into a single cell suspension and labeled with 1μM CellTrace Far Red (ThermoFisher Sci). Targets were then divided into two populations: NS4b_2488_ targets were labeled with Cell Trace Violet (ThermoFisher Sci) 5mM for 10^−6^M peptide loaded targets and 0.5mM for non-loaded negative control) and ENV_641_ targets were labeled with CFSE (ThermoFisher Sci) 5mM for 10^−6^M peptide loaded targets and 0.5mM for non-loaded negative control. All labeling was done as described in manufacturer instructions. Peptide loading was 1 hour at 37°C. After thorough washing, targets were pooled and injected intravenously (i.v.) into infected and naïve mice. After 4 hours mice were sacrificed spleen disassociated to a single cell suspension. Splenocytes were gated on total targets (CellTrace Far Red positive) then CellTrace Violet high and low or CFSE high and low. Calculation of killing: 100% - [(% peptide bearing targets IMMUNE recipients / % no peptide targets IMMUNE recipients) / (% peptide bearing targets NAIVE recipients / % no peptide targets NAIVE recipients) x 100].

## STATISTICS

GraphPad Prism^®^ version 9.2.0 (GraphPad Software) was used for data analysis. All experiments were performed at least twice. Significance was determined as described in figure legends.

## ACKNOWLEDGEMENTS

Authors wish to thank Drs Jason Netland and Michael J. Bevan for sharing the WNV-I TCR transgenic mice; the NIH tetramer facility for providing Class I MHC tetramers; Dr. Deepta Bhattacharya for providing reagents; Dr. Anne Wertheimer for assistance with human sample processing; and Drs. Nancy R. Manley, Lauren I. R. Ehrlich, Bonnie J. LaFleur, Gregory Sempowski, and Laura Hale for their critical reading of the manuscript and suggestions. Graphics were created in Biorender.com.

## AUTHOR CONTRIBUTIONS

J.L.U., J.N-Ž., J.D. and M.R.M.vdB. designed the research; J.L.U., M.J., C.M.B., S.S., and C.P.C. performed experiments and analyzed the data; K.O.M., M.C.L., M.P.B. contributed human samples; J.L.U. prepared the figures; J.L.U. and J.N-Ž. wrote the manuscript; all authors edited the manuscript.

## DATA AVAILABILITY

All data available upon request.

