## Supplementary Figures for "Quantitative Restoration of Immune Defense in Old Animals Determined by Naïve Antigen-Specific CD8 T cell Numbers"

Supp. Figure 1

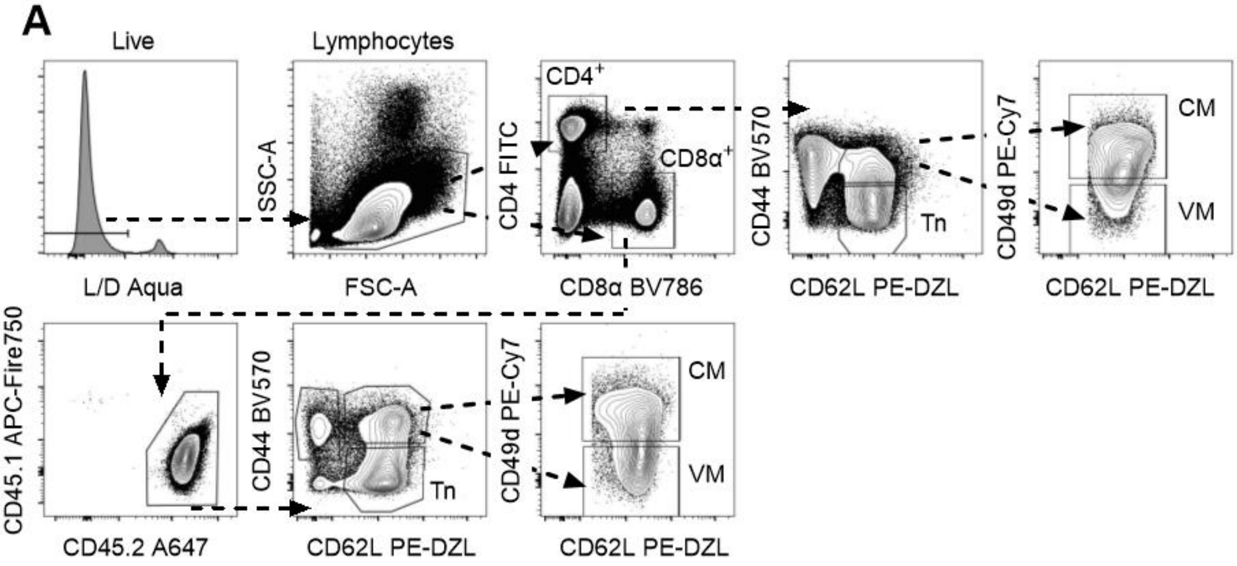


Supplemental Figure 1

**Representative FCM gating strategy for phenotypes in Figure 3.**

### Supp. Figure 2

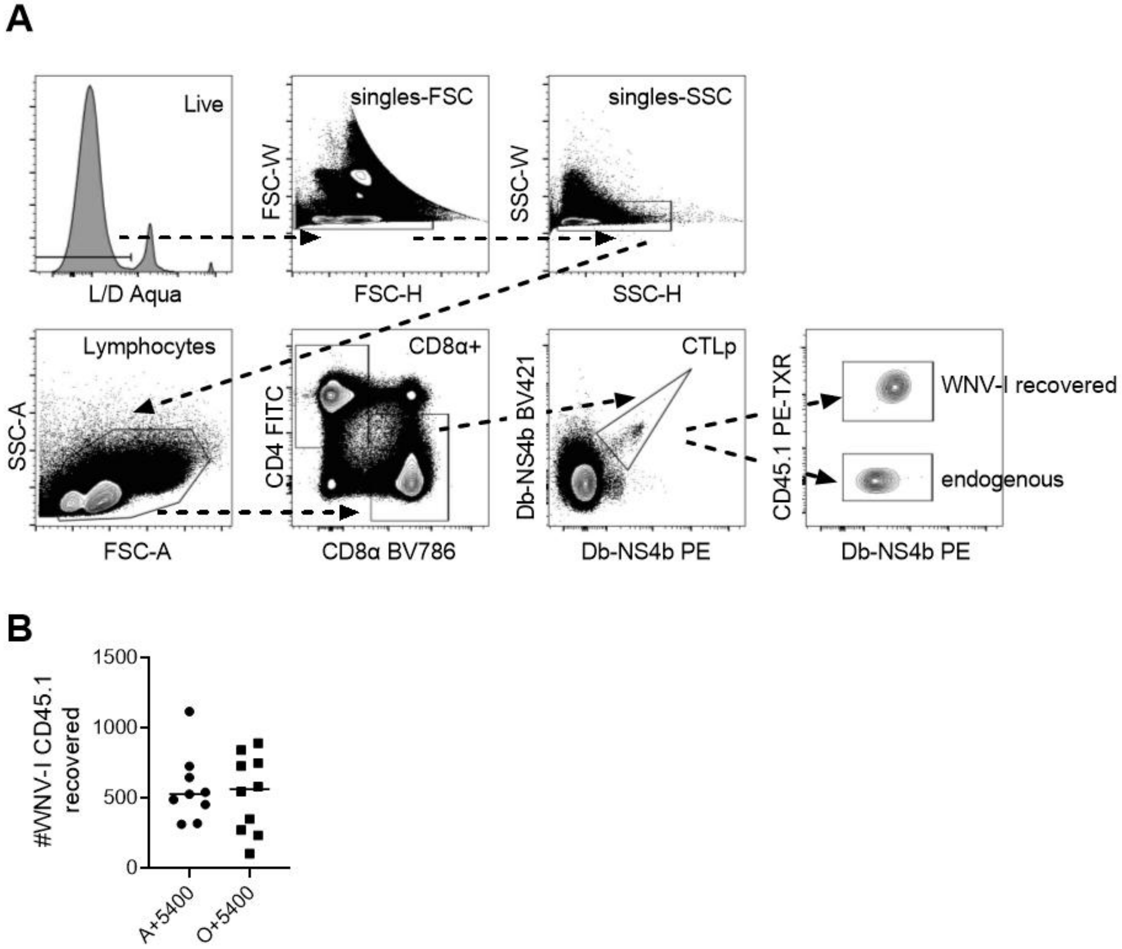


Supplemental Figure 2

**Equal engraftment of WNV-I CD8 T cells 24 hours post-transfer into adult and old.** (A) Representative gating strategy for the enumeration of endogenous (CD45.2) or donor (CD45.1) WNV-specific CD8 T cells. (B) 5400 WNV-I CD8 T cells were transferred into adult (3-5 months) and old (19-22 months) mice to evaluate engraftment into SLO. At 24 hours post-transfer, the number of CD45.1 NS4b-Tg CD8 T cells were enumerated from pooled spleen and LNs alongside endogenous CTLp shown in Figure 3K.

Supplemental Table I

Demographics of human subjects, Figure 1.

| | Asymptomatic | WNF | WNM | WNME |
|---|---|---|---|---|
| Total | n=20 | n=15 | n=12 | n=18 |
| Age<br>(years ±SEM) | 50.09 ± 2.865 | 64.63 ± 4.586 | 59.71 ± 3.037 | 69.31 ± 2.834 |
| sex | n=10 male<br>n=10 female | n=10 male<br>n=5 female | n=5 male<br>n=7 female | n=13 male<br>n=5 female |
| days post-onset (±SEM) | N/A | 629.5 ± 119 | 1076 ± 211 | 955.1 ± 108 |
